# Tumour-Associated Extracellular Vesicles Carry Extrachromosomal DNA and Influence Recipient Cell Phenotypes

**DOI:** 10.64898/2026.11.24.769016

**Authors:** Ensieh M. Poursani, Behdad Delavari, Vu V. Pham, Bleron Zajmi, Anahid Ehteda, Jake Baum, Orazio Vittorio

## Abstract

Extrachromosomal circular DNA (ecDNA) is a key driver of oncogene amplification, tumour evolution, and therapeutic resistance.

Extracellular vesicles (EVs) are nanoscale membrane bound structures released by cells into the extracellular space. These vesicles play important roles in modulating tumor microenvironment and immune responses via transferring biomolecular cargos between donor and recipient cells. Although recent studies have reported the presence of DNA in EVs, whether extracellular vesicles (EVs) can transport and spread extrachromosomal circular DNA (ecDNA) remains unknown.

Here, we first investigated whether tumor-derived EVs can carry ecDNA and then, explored its influence on behaviours of recipient cells. In brief, EVs were isolated from the ecDNA-positive colorectal cancer cell line COLO320DM and characterised by nanoparticle tracking analysis, their identity was confirmed by the expression of the EV markers CD81 and Alix. The presence of genomic DNA in the EV fractions was analysed by PCR. Confocal microscopy further demonstrated the uptake of EV-associated DNA by recipient cells. Importantly, long-read Oxford Nanopore sequencing analysis of EV-associated DNA identified MYC-containing ecDNA, providing evidence of a link between oncogenic ecDNA and tumour-derived EVs. Functionally, prolonged exposure to COLO320DM-derived EVs reduced surface HLA-ABC expression in HL60 cells, suggesting that EVs may affect antigen presentation and immune recognition. In HeLa cells, treatment with COLO320DM-derived EVs increased cell mobility and migration. Notably, in colony formation assays, exposed HeLa cells did not form discrete colonies; instead, they showed a dispersed growth pattern extending into the surrounding area. In sum, these findings provide evidence that tumor-derived EVs carry oncogenic ecDNA, and suggest tumor-derived EVs may contribute to cell-cell communication and influence recipient cell behaviour, highlighting a potential role for EV-associated ecDNA in cancer progression.

## INTRODUCTION

Extracellular vesicles (EV) are small membrane-bound particles that transfer different types of molecular cargo, including mRNA, microRNA, long non-coding RNA (lncRNA), DNA, proteins, and lipids between cells. EVs are typically enriched in protein markers such as CD63, CD9, and CD81^1^.They can enter into other cells by merging their membrane with recipient cells plasma membranes. This enables the transfer of molecular and genetic cargo between cells and plays a vital role in mediating intercellular communication ^2,3,4^.

Tumour-derived EVs contribute to a variety of cancer-related biological processes such as cell proliferation, migration, invasion, angiogenesis and interaction with tumour microenvironment. Furthermore, tumour-derived EVs can modulate immune responses and subsequently contribute to tumor progression and therapy resistance^5^.

The immune system interacts continuously with tumors throughout their development and metastatic spread. This dynamic and complex communication can support tumor growth and is now recognised as a fundamental hallmark of cancer. ^6,7^.

Immune evasion is the process by which pathogens or cancer cells avoid detection and destruction by the host’s immune system, allowing them to persist and grow. Cancer cells can develop sophisticated mechanisms to escape immune system. This also contributes to their resistance to certain treatments. ^7,8^.

Malignant cells use different strategies to avoid immune recognition by disrupting the normal processes of antigen processing and presentation. These strategies include modifying surface antigens, producing immunosuppressive molecules, and affecting the activity of immune cells^7^.

ecDNA was first detected in cancer in 1965^9^. They are a well-recognised origins for amplified oncogenes in human cancers^10,11^. They are typically from hundreds of kilobases to several megabases^12,13^. EcDNAs do not have centromere and segregate randomly during cell division^14^. Their chromatin is less compacted compared to the chromosomes which makes them susceptible to high rates of mutations^15^. Therefore, they can promote genetic heterogeneity, therapy resistance, and rapid genome alterations that may consequently accelerate disease progression^15,16^. While the contribution of ecDNA to tumor evolution and therapy resistance is now well recognised, the mechanisms underlying these effects are still poorly defined.

Given that previous studies have reported that EVs can carry DNA and oncogenic genetic materials between cells, we propose that tumour-derived EVs may also carry oncogenic extrachromosomal circular DNAs (ecDNAs) to extracellular environment.

Oncogenic ecDNAs can potentially be exchanged between cancer cells and immune cells. If these ecDNAs and their products can be integrated into immune cells, they may impair the immune system’s ability to recognize malignant cells as threats.

In this study, we investigated the presence of ecDNA in EVs derived from ecDNA-positive colorectal cancer cell line COLO320DM. We extracted DNA from EVs and analysed long-read Nanopore sequencing to identify oncogenic ecDNA. We further investigated whether exposure to these EVs alters phenotypic characteristics relevant to tumour progression, including HLA-ABC surface expression, cell mobility, migration, and colony-forming behaviour.

Our findings provide evidence that tumour-derived EVs carry oncogenic ecDNA. Furthermore, since exposure to these EVs may alter characteristics of recipient cells, exposure may represent a previously underappreciated mechanism linking horizontal genetic communication with tumour progression, immune evasion, and metastatic behaviour. This study highlights that oncogenic ecDNAs can potentially be transferred using tumour-derived EVs and subsequently contribute to cancer progression and immune evasion.

## RESULTS

### COLO320DM-Derived EVs Are Taken Up by Recipient Cells

We first examined whether COLO320DM cells can release EVs into the culture medium and whether they can be taken up by other cells. Using transmission electron microscopy (TEM), we observed vesicle-like structures of EVs ranging from 60 to 330 nm (Fig. 1a).

**Figure 1.**
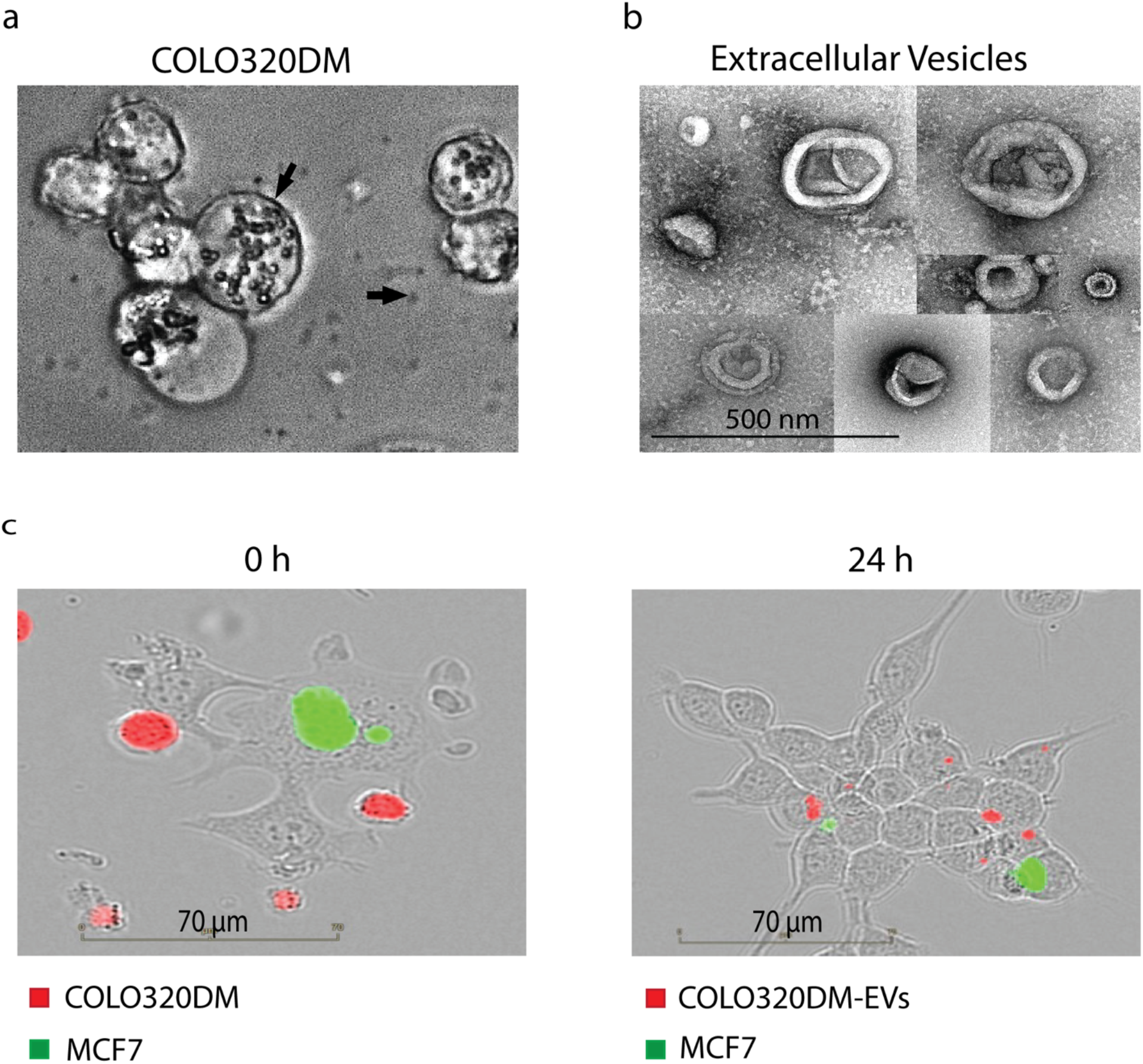
Characterisation of COLO320DM-derived extracellular vesicles (EVs) and their transfer to recipient cells. (a) Bright-field microscopy images indicating EV-like structures (black arrows) on the COLO320DM cell surface and in the supernatant (40X magnification). (b) Transmission electron microscopy (TEM) images of EVs isolated from COLO320DM-conditioned medium using field-flow electrophoresis (FFE), followed by concentration through ultracentrifugation. (c) Co-culture of COLO320DM cells (labelled with red Bodipy dye) and GFP-expressing MCF7 cells, showing the transfer of COLO320DM-derived fluorescent material to recipient MCF7 cells.

To find whether these vesicles could enter recipient cells, EVs were isolated from COLO320DM culture supernatants by FFE and subsequent ultracentrifuge to concentrate them. Then, EVs were resuspended in PBS and labelled with BODIPY fluorescent dye for 20 minutes. After three washes with PBS to remove excess dye, the labelled EVs were added to pre-seeded MCF7 cells stably expressing green fluorescent protein (GFP). Live-cell images were captured every 4 hours using the IncuCyte SX5 system.

We observed red fluorescence signal within the cytoplasm of GFP-expressing MCF7 cells 24 hours following exposure with red labelled-COLO320DM cells (Fig. 1b). This suggests that material associated with COLO320DM cells can enter recipient cells. Overall, these findings support the ability of tumour-derived EVs to interact with and potentially transfer their contents to other cells. This highlights their possible role in intercellular communication.

### COLO320D-derived EVs contain genomic DNA sequences

To investigate whether EVs contain genomic DNA, ultracentrifuge was used to isolate EVs from COLO320DM cell supernatants and DNA was extracted using an exoDNA extraction kit. PCR was applied with MYC gene primers to confirm the presence of genomic DNA in the isolated EVs (Fig. 2a).

**Figure 2.**
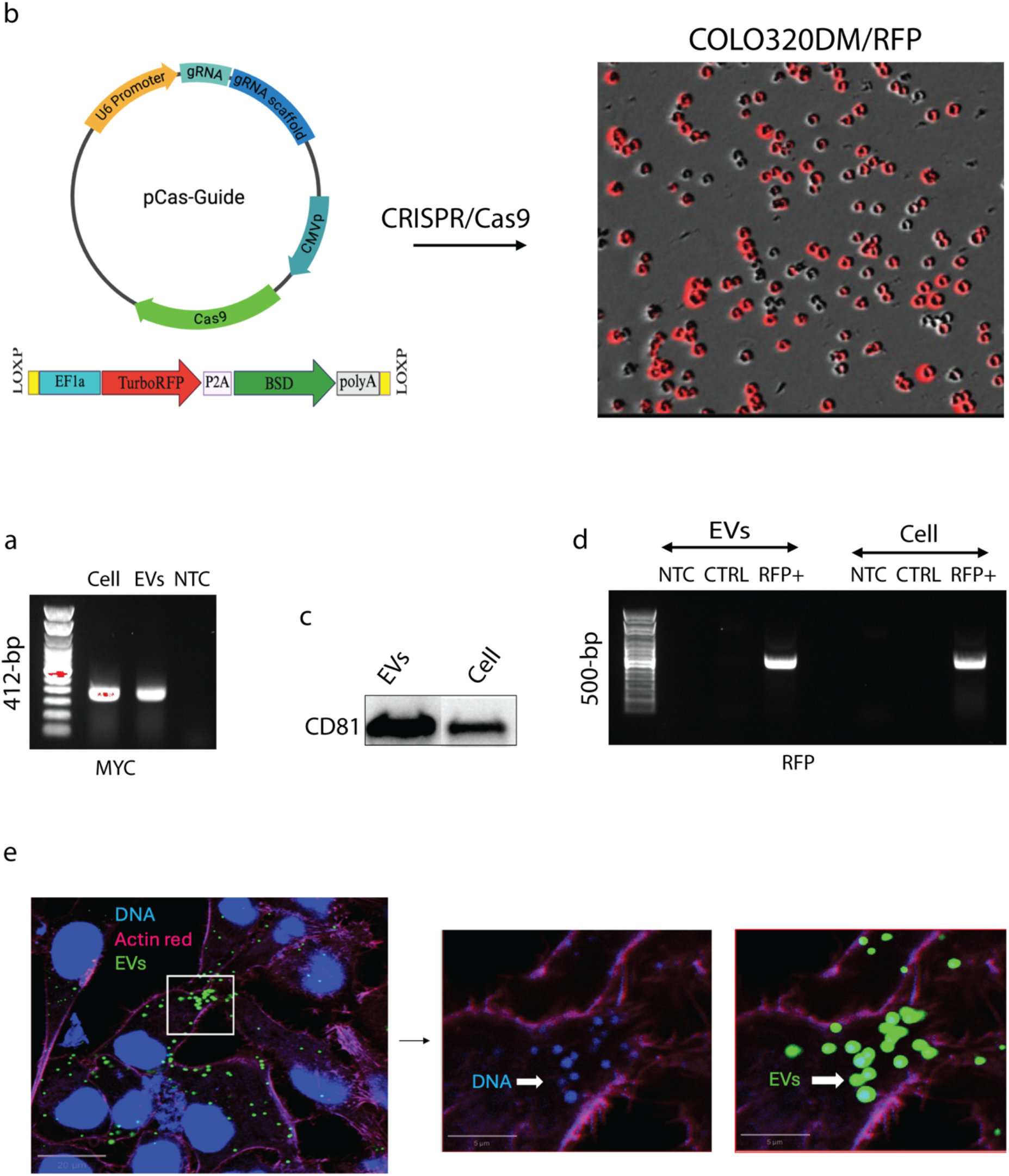
Detection of genomic DNA in COLO320DM-derived EVs and assessment of EV uptake by recipient cells. (a) PCR detection of the MYC gene in genomic DNA and EV-derived DNA. (b) Generation of a stable COLO320DM/RFP cell line expressing turboRFP following CRISPR–Cas9-mediated knock-in of a turboRFP–blasticidin resistance cassette into the downstream of MYC gene. (c) Western blot analysis of the EV-associated marker of CD81 in whole-cell lysates and isolated EV preparations. (d) PCR detection of turboRFP sequences in genomic DNA of COLO320DM, COLO320DM/RFP cells and their derived EVs. (e) Representative confocal microscopy images showing the uptake of BODIPY-labelled COLO320DM-derived EVs by HeLa recipient cells. DAPI staining revealed DNA-associated fluorescent signals overlapping with BODIPY-labelled EV signals.

Moreover, CRISPR-Cas9-mediated genome editing was used to insert a donor DNA encoding turboRFP (donor DNA; *EF1a promoter-turboRFP-P2A-Blasticidin-polyA*) into the 3’ terminus of the MYC gene on both chromosomal and extrachromosomal DNA (ecDNA) and stable COLO320DM/RFP cell line was selected using Blasticidin selection following by cell sorter (FACS) (Fig. 2b).

To confirm the integration of RFP into the genomic DNA, first EVs were tested with western blotting for CD81 (Fig. 2c). Then, specific primers amplifying genomic RFP were used. PCR analysis revealed the presence of RFP genomic DNA in both COLO320DM/RFP cells and their derived EVs, while no signal was detected in parental COLO320DM cells and its EVs (Fig. 2d).

Furthermore, COLO320DM-derived EVs were labelled with green Bodipy dye and subsequently added to HeLa cells for an overnight to assess EV uptake. Next day, Hela cells were fixed, stained with actin red (phalloidin) to visualize the cytoskeleton, and mounted using DAPI-mounting medium to label the nuclei. Confocal microscopy was used to analyse samples using z-stack imaging. Confocal microscopy images confirmed the uptake of labelled EVs by HeLa cells. Of note, we observed a degree of co-localisation in the green Bodipy and DAPI fluorescent signals, suggesting a potential association between EVs and DNA (Fig. 2e). However, additional experiments would be required to confirm this co-localization and the presence of DNA within EVs.

In addition, confocal microscopy revealed signals suggestive of DNA-containing cargo being released from the nuclei of both COLO320DM and HeLa cells (Fig. 3). These observations raise the possibility that cells may directly package nuclear DNA into small vesicles and subsequently release it into the extracellular environment. This process could potentially contribute to intercellular communication and the transfer of genetic material between cells. However, further investigation is required to confirm the nuclear origin of the DNA, verify its presence within these vesicles, and elucidate the mechanisms underlying their formation and release.

**Figure 3.**
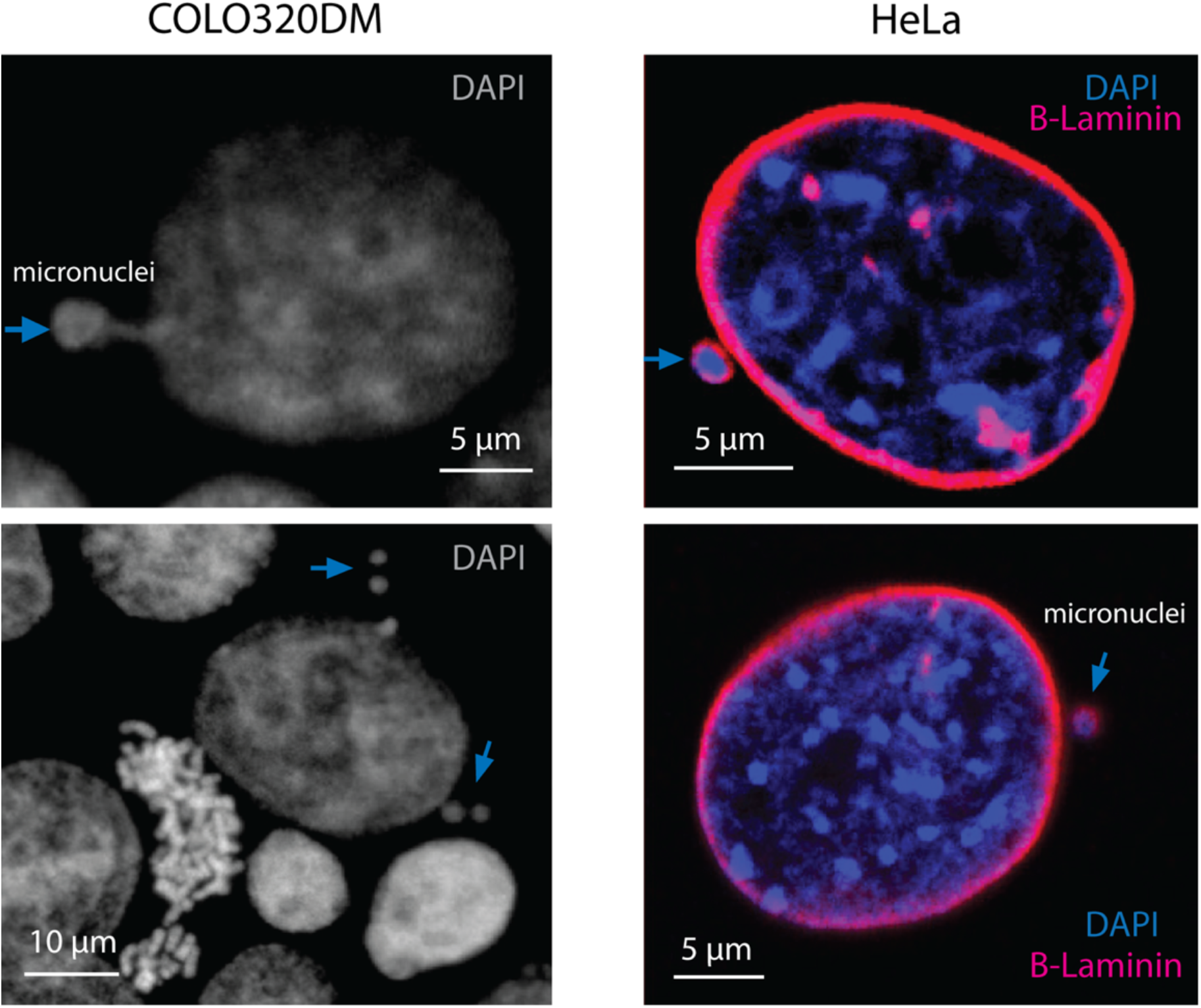
Detection of micronuclei in COLO320DM and HeLa cells. COLO320DM cells were stained with DAPI, while HeLa cells were co-stained with DAPI and β-laminin to visualise nuclear envelope.

### Nanopore sequencing identifies circular DNA signatures in COLO320DM-derived EV-associated DNA

To investigate whether COLO320DM-derived EVs are associated with extrachromosomal circular DNA (ecDNA), we performed a series of experiments using EVs isolated from COLO320DM cell culture supernatants.

Briefly, the supernatant from COLO320DM cell cultures was collected, filtered, and centrifuged to remove cells and cell debris, as described in material and method section. Then, the supernatant was fractioned using Free Flow Electrophoresis (FFE) to separate extracellular particles according to their electrophoretic mobility (Fig. 4a). The FFE isolated fractioned were initially analysed using dot blot with CD81 antibody (Fig. 4b) and ZetaView (Fig. 4c-d) to detect EV-enriched fractions.

**Figure 4.**
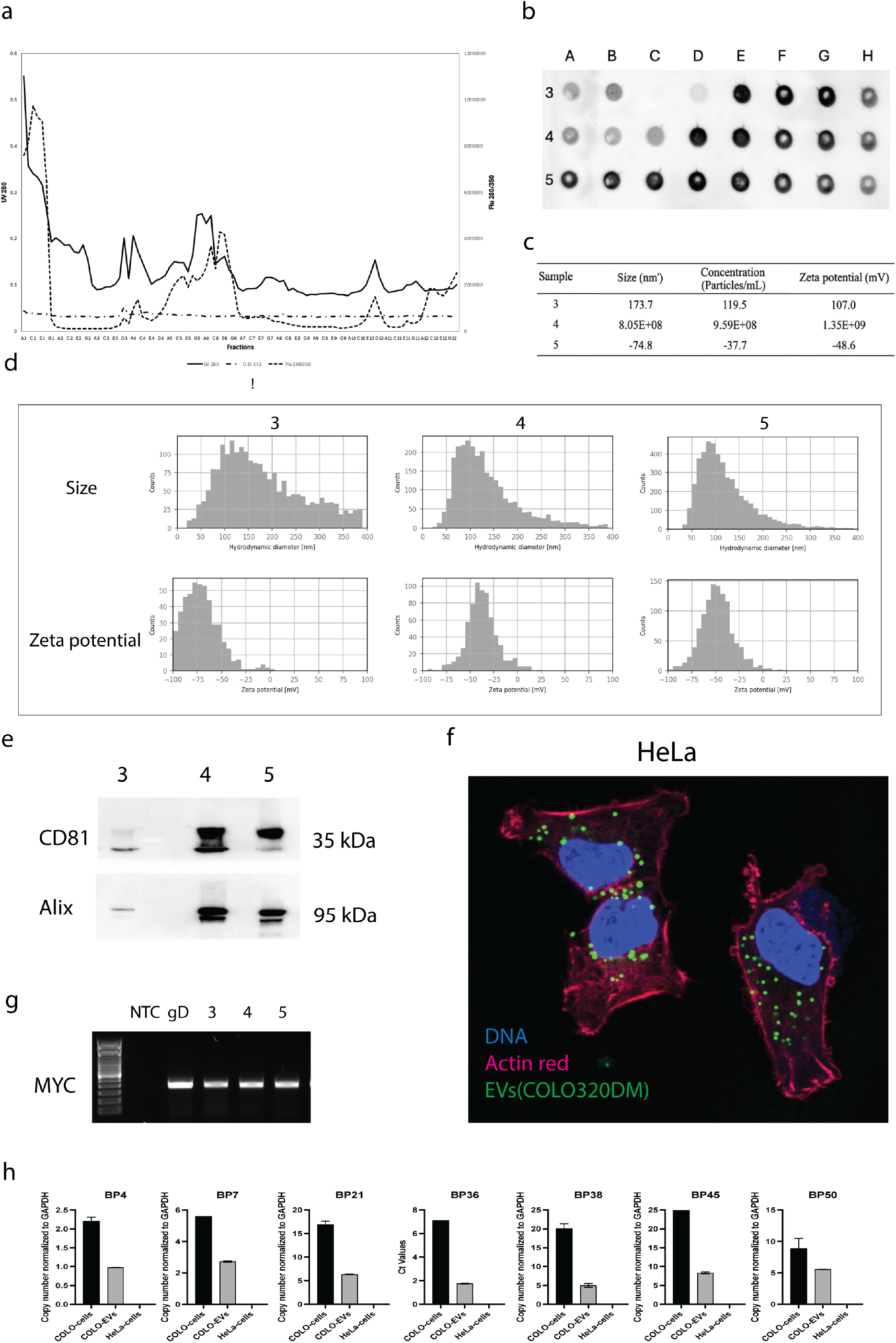
Free flow electrophoresis (FFE) fractionation and characterisation of extracellular vesicles (EVs).(a) FFE separation profile showing UV absorbance at 280 nm (UV280), conductivity and fluorescence intensity across the collected fractions. (b) Dot blot of the collected FFE fractions using an anti-CD81 antibody to identify the EV containing fractions. Positive CD81 signal was seen in the fractions corresponding to rows 3-5. (c & d) Characterisation of the CD81 positive fractions by ZetaView nanoparticle tracking analysis (NTA) to determine particle size, concentration and zeta potential. This provides a characterisation of the EV enriched fractions (rows 3, 4 and 5) following FFE separation. (e) Western blot analysis of EV fractions showing the expression of the EV-associated markers of CD81 and ALIX. (f) Confocal microscopy images showing the uptake of COLO320DM-derived EVs by HeLa cells. (g) PCR detection of MYC DNA in the EV fractions. Lanes 3–5 represent FFE-separated EV fractions, gD is COLO320DM genomic DNA as a positive control, and NTC is non-template negative control of PCR. (h) Real-Time PCR analysis of ecDNA Breakpoints in COLO320DM-associated EVs. Specific primer sets were used for amplifying seven distinct breakpoints (BPs) in COLO320DM-derived ecDNA to investigate the presence of ecDNA sequences in extracellular vesicles (EVs) released by COLO320DM cells. Genomic DNA isolated from COLO320DM cells was used as a positive control, and genomic DNA from HeLa cells used as a negative control. Copy numbers of each breakpoint were normalized to GAPDH levels.

After characterizing FFE-isolated fractions they were divided into three groups (groups 3, 4 & 5), and each group was separately examined by transmission electron microscopy TEM. Image analysis revealed vesicle-like structures ranging in size from approximately 60 to 200 nm (Fig. 1).

Next, western blotting for CD81 and Alix was used to confirm the presence of EV-associated markers (Fig. 4e).

Selected fractions were also used for labelling with fluorescent dye followed by confocal microscopy, as well as downstream DNA analysis (Fig. 4f).

Upon PCR validating of the MYC genomic DNA in EV-associated fractions (Fig. 4g), the EV-DNA was subjected to long-read Nanopore sequencing to characterize complex DNA structures.

Sequencing data were subsequently analysed using the CoRAL (Complete Reconstruction of Amplifications with Long Reads), which is specialized to reconstruct circular amplicon architectures from long-read sequencing data (Fig. 5a). The predicted amplicon structures showed similarities to the ecDNA structures previously characterized in COLO320DM genomic DNA (Fig. 5b). These findings were further supported by real-time PCR analysis targeting specific COLO320DM ecDNA breakpoints (BPs), which detected these BP sequences in COLO320DM-associated EVs (Fig. 4h), providing more evidence that EV-associated DNA may contain ecDNA-derived sequences.

**Figure 5.**
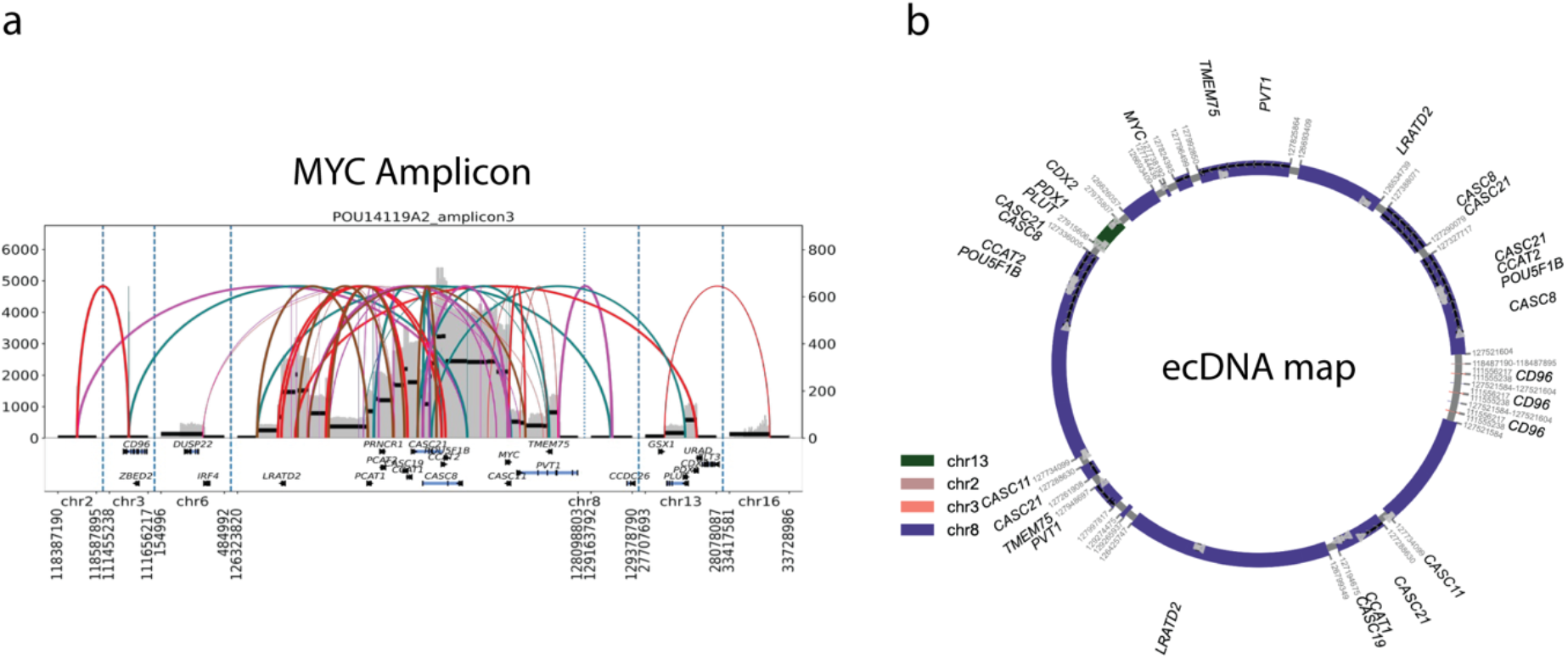
Reconstruction of ecDNA from EV-derived DNA using long-read nanopore sequencing. (a) Reconstruction of MYC-containing ecDNA from EV-DNA by analysing nanopore sequencing data with CoRAL, showing the predicted amplicon structure (a) and circular ecDNA map (b).

Together, these results suggest that DNA associated with COLO320DM-derived EVs may contain sequences originating from cellular ecDNA.

### EVs derived from COLO320DM cells reduce HLA-ABC on the HL60 cell surface

To explore whether EVs released by ecDNA-positive COLO320DM cell line could affect HLA-ABC expression in the immune cells, HL60 leukemia cells were treated with COLO320DM-derived EVs for four weeks. Then, both HL60-treated and untreated control cells were stained with an anti-HLA-ABC antibody and propidium iodide (PI), followed by analysis by a Flow Cytometer. The results showed about 10% reduction in mean HLA-ABC fluorescence intensity in EV-treated cells compared with the untreated controls, suggesting that prolonged exposure with COLO320DM-EVs may affect antigen presentation and immune recognition by reducing HLA-ABC on cell surface (Fig. 6).

**Figure 6.**
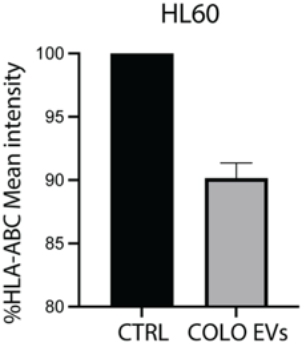
Flow Cytometric Analysis of HLA-ABC Expression on HL60 Cells following Treatment with COLO320DM-associated EVs. HL60 human leukemia cells were treated with COLO320DM-derived EVs for four weeks. Surface expression of HLA-ABC molecules was assessed by flow cytometry in treated (COLO EVs) and non-treated (CTRL) samples.

### COLO320DM-derived EVs induce cell migration in Hela cells

To investigate the effect of COLO320DM-derived EVs on HeLa cell migration, we first performed wound-healing assay. As shown in Fig. 7, treating cells with COLO320DM-derived EVs significantly increased cell mobility, compared with the untreated control group. This finding was then confirmed by a Transwell migration assay. Which showed a higher migration rate (64.87%) in the EV-treated group compared with the control group (38.4%), suggesting that COLO320DM-derived EVs enhance the migratory capacity of HeLa cells (Fig. 7a-b).

**Figure 7.**
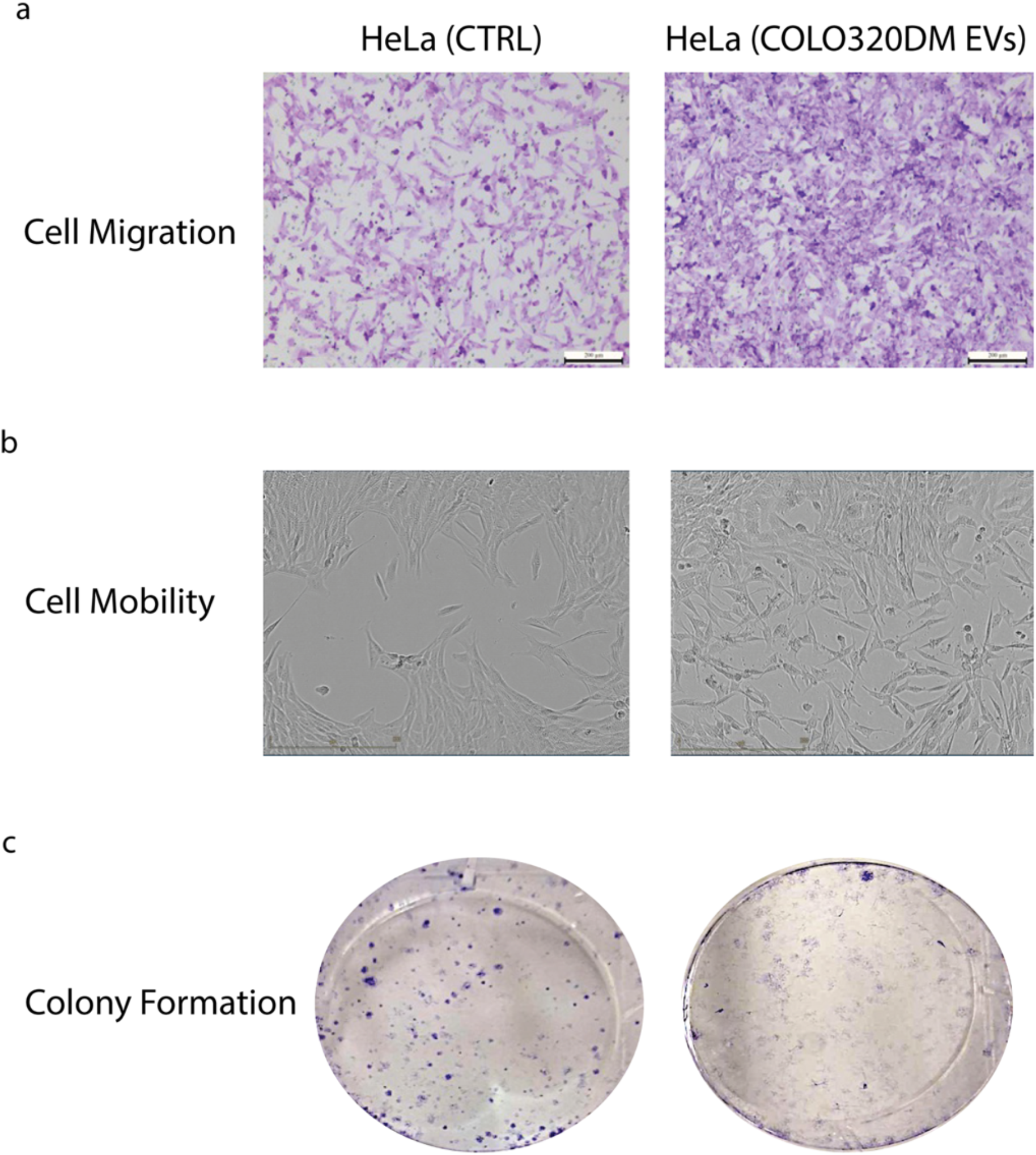
Functional assays analysing the effects of COLO320DM-derived EVs on the HeLa cell characteristics. (a) Transwell assay indicating the migratory capacity of HeLa cells following treatment with COLO320DM-derived EVs. (b) Wound-healing assay showing increased mobility activity of HeLa cells after EV treatment. (c) Colony formation assay demonstrating changes in HeLa cell colony morphology, with forming more dispersed colonies in EV-treated cells compared to the non-treated control. Data are representative of at least three independent experiments.

### COLO320DM-derived EVs inhibit colony formation and induced dispersed growth pattern in Hela cells

HeLa cells were exposed to COLO320DM-derived EVs for two weeks. Then, cells were stained with Crystal Violet to assess colony formation. EV-treated cells displayed fewer colonies compared with the untreated cells. In addition, instead of growing as well-organized compact colonies, the treated cells scattered and grew dispersedly (Fig. 7c). These findings suggest that COLO320DM-derived EVs may affect both the colony-forming capacity and growth behaviour of HeLa cells.

## DISCUSSION

Extracellular vesicles (EVs) contribute to cell-cell communications. They can play an important role in cancer progression, extracellular matrix remodelling, metastasis and immune system suppression^17^ by transferring biomolecules such as lipids, proteins, RNA and DNA between cells. Previous studies have demonstrated that tumour-derived EVs can carry DNA cargo (10 to 100kb) which highlights their potential role in cancer development ^18^. However, whether EVs can carry extrachromosomal circular DNA (ecDNA) remain poorly understood.

In this study, we provide evidence that EVs derived from COLO320DM cells contain genomic DNA, including MYC-containing circular DNA, consistent with COLO320DM genomic ecDNA. Using different molecular analysis including PCR, long-read nanopore DNA sequencing and CoRAL, we detected circular MYC-containing DNA in the EV-derived DNA. These findings propose that EVs may contribute to dissemination of oncogenic DNA within tumour microenvironment.

We also explored the potential effects of COLO320DM-derived EVs on responses of cancer and immune cell models. We found that COLO320DM-derived EVs increased HeLa cell mobility and migration in wound-healing and Transwell migration assays. Furthermore, EV-treated HeLa cells exhibited altered colony formation characteristics. EV-treated cells had fewer colonies and a more dispersed growth pattern compared to the non-treated cells. These findings indicate that tumour-derived EVs can affect the migratory behaviour and growth pattern of recipient cells. However, whether these effects are directly mediated by ecDNA or by other EV-associated components remains unclear and needs more studies.

We also investigated the potential effect of tumour-derived EVs on immune-related responses of recipient cells. prolonged exposure of HL60 cells to COLO320DM-derived EVs resulted in a approximately 10% reduction in surface HLA-ABC expression. HLA-ABC molecules are essential for presenting tumour-derived antigens to immune cells. Therefore, changes in their surface expression level may influence immune recognition. However, further functional studies are needed to confirm the contribution of tumor-derived EVs on immune recognition.

In conclusion, this study provide evidence that tumour-derived EV s can carry circular oncogenic DNA including ecDNA. Also, exposure to these EVs is associated with changing migratory characteristics, colony formation and HLA-ABC expression in the recipient cells.These findings suggest a potential link between EV-mediated oncogenic DNA dissemination and tumour progression. However, further studies are needed to determine whether transferred ecDNA remains functional or whether other EV-derived components are responsible for the observed effects.

## METHODS

### Cell culture

Human colon cancer cell line COLO320DM cell line was kindly provided by Prof. John Mariadason from Olivia-Newton John Cancer Research Institute (Australia) and cultured in DMEM with 10% fetal bovine serum (FBS).

HL-60 and HeLa cells were purchased from ATCC and cultured in DMEM with 10% FBS.

### COLO320DM stable transfection

A donor DNA construct containing turboRFP and a Blasticidin resistance gene (KN2.0 linear donor [RFP-BSD], Cat# CW310646, OriGene) was knocked-in into the COLO320DM cells using CRISPR-Cas9 with two guide RNAs targeting the 3′ untranslated region (3′-UTR) of MYC oncogene (gRNA1: AACACAAACTTGAACAGCTA, gRNA2: TCAAATGCAACCTCACAACCT). Transfection was performed using Lipofectamine 3000 according to the manufacturer’s instructions. Two weeks post-transfection, cells were initially selected using Blasticidin (10µg/ml), followed by fluorescence-activated cell sorting (BD FACSAria Fusion) to isolate turboRFP-positive cells.

### Cell migration assay

Cell migration was assessed using 12-well chambers (8-μm pore size) according to the manufacturer’s protocol (Cat# 353182, Corning). Briefly, HeLa cells (3 × 10^4^ cells/well) were seeded in the top chamber of the inserts containing low serum Media (2% FBS) and COLO320DM-derived EVs. The bottom chamber included media with chemoattractant (20% FBS) and plates were incubated in a humidified incubator with 5% CO2 at 37 °C for 16 h. At the endpoint, transwell inserts were stained with crystal violet and imaged. The number of migrated cells was quantified using ImageJ software.

### Wound-healing assay

HeLa cells (2 × 10^4^ cells/well) were seeded in 96-well plates and cultured overnight to reach 90% confluency. The next day, a gap was generated on a confluent monolayer of cells using an Incucyte® 96-well Woundmaker Tool (Satorious, Germany) and cells were washed with PBS. Cells were then treated with COLO320DM-derived EVs. After treatment plates were immediately incubated at 37°C, 5% CO_2_. Cells were imaged every 4 h. Images and data were obtained using IncuCyteS5 (Essen Bioscience) with a 10 × object. Incucyte® Scratch Wound Analysis Software Module (Cat. No. 9600–0012) was used to automatically measure the scratch length.

### EVs isolation by Ultracentrifuge

COLO320DM cells were cultured in DMEM (serum-free media) in a T175 flask for 48 hours. Cell supernatant was collected and centrifuged at 1500 rpm for 5 minutes to pellet the cells. Next, the supernatant was transferred to a clean tube and centrifuged at 3,000 rpm for 10 minutes to pellet residual cells. The supernatant was then filtered through a 0.45 µm filter. Then, the supernatant was ultracentrifuged at 100,000 g for 2 hours at 4 °c. The supernatant was carefully removed, and EVs were resuspended in 200 µl of sterile PBS.

### EVs separation using Free-flow electrophoresis (FFE)

Cells were incubated overnight in T175 flasks containing serum-free Dulbecco’s Modified Eagle Medium (DMEM). Conditioned medium was collected and centrifuged sequentially at 1,500 rpm for 5 min, 3,000 rpm for 10 min, and 10,500 rpm for 1 h at 4°C to remove cells, cellular debris, and larger particles. The supernatant was then filtered through a 0.45 µm membrane prior to separation by free-flow electrophoresis (FFE).

FFE separation was conducted according to the method described by Preußer et al. (2022)^19^, with modifications. The clarified supernatant was introduced into the FFE separation chamber, where a continuous laminar flow of separation buffers transported the sample through an electric field oriented perpendicular to the flow direction. This process separated particles based on differences in electrophoretic mobility, determined primarily by surface charge and the pH of the separation medium. The separation buffers maintained a controlled electrophoretic environment, enabling particles with distinct charge characteristics to migrate along specific trajectories. After separation, fractions were collected from multiple outlets and analysed to identify EV-enriched populations.

Fractions were screened for the extracellular vesicle (EV) marker CD81 using dot blot analysis. CD81-positive fractions corresponding to rows 3-5 were subsequently characterised by ZetaView nanoparticle tracking analysis (NTA) to determine particle concentration, hydrodynamic size distribution, and zeta potential.

### Western Blot

EVs were lysed with RIPA lysis buffer supplemented with protease and phosphatase inhibitors (1x cOmplete Mini, EDTA-free Protease Inhibitor Cocktail, and PhosSTOP™ phosphatase inhibitor, Sigma-Aldrich, Schnelldorf, Germany). BCA Protein Assay Kit (Thermo Fisher Scientific, USA) was used to measure protein concentration. An equal amount of protein from EVs and whole cell lysate was loaded onto 4–20% SDS-PAGE gels (Mini-PROTEAN TGX Gels, Bio-Rad, USA) and transferred to the nitrocellulose membranes (Amersham™ Protran® 0.45 µM, GE Healthcare, Germany). Membranes were blocked in 5% BSA/TBS-T (1X TBS + 0.1% Tween-20) for 1 h and probed with antibodies against CD81 and Alix proteins for an overnight at 4 °C. Then membrane was washed in TBS-T and incubated with anti-rabbit (1:5000) or anti-mouse (1:5000) IgG secondary antibodies for 1 h at room temperature. The resultant signal was visualized using Clarity ™ Western ECL Substrate (Bio-Rad, USA).

### DNA extraction

COLO320DM-derived EVs were treated with DNase I and DNA was extracted using EasySC Exosome DNA Purification Kit (Amerigo, Catalog Number: D035-1, D035-2), based on manufacturer’s instruction. Whole genomic DNA was extracted from COLO320DM cells using QIAGEN DNeasy Blood & Tissue kit, based on the manufacturer’s instruction. DNA purity and concentration were assessed using Nanodrop 2000 (Thermo Fisher Scientific, USA) and QC was done using Qubit.

### PCR

To assess the presence of MYC and turboRFP genes in both cellular and EV-DNA, PCR was performed using a Taq 2× Master Mix (NEB, Catalog Number: M0270). The primer sequences used were as follows:

MYC-F: 5′-GGACTTGTTGCGGAAACGACGAG-3′, MYC-R: 5′-CTATATACTAGGATTGAAATTCTGTGTAACTGC-3′, turboRFP-F: 5′-ATGAGCGAGCTGATCAAGGAGAAC-3′, turboRFP-R: 5′-TCTGTGCCCCAGTTTGCTAGGG-3′. Each PCR reaction was set up in a total volume of 25 µL using 100 ng of DNA template extracted from cells or EVs. Thermal cycling was as follows: initial denaturation at 95 °C for 2 minutes; 30 cycles of denaturation at 95 °C for 30 seconds, 60 seconds annealing at 56 °C for cMYC and 58°C for turboRFP, extension at 68 °C for 20 seconds; followed by a final extension at 68 °C for 4 minutes. PCR products (8 µL) were analyzed by agarose gel electrophoresis.

### qPCR-Based Quantification of ecDNA Breakpoint Copy Numbers

Quantitative real-time PCR (qPCR) was performed to assess the relative abundance of COLO320DM-associated ecDNA breakpoint (BP) sequences in EV-derived DNA, COLO320DM genomic DNA and HeLa genomic DNA. Target-specific primer sets were designed to amplify distinct ecDNA breakpoints.

PCR reactions were prepared using SYBR Green PCR master mix, with 50 ng genomic DNA as instructed by manufacturer.

PCR amplification was performed on Bio-Rad Real-Time PCR CFX96 using the following cycling conditions: initial denaturation at 95 °C for 3 min, followed by 33 cycles of 95 °C for 10s and 59 °C for 30s.

Copy number variation was assessed by quantitative real-time PCR (qPCR) by comparing the cycle threshold (*C_t_*) values of the target gene

Primer sequences amplifying BPs are as follow:

BP4-F; AAAGTGGCTATGTTCTGTCC, BP4-R; CCATCCTACATTTCACATGG, BP7-F; CCCTTTGTCAGATGAGTAG, BP7-R; GCTGTGCATGTATATCACTTC, BP21-F; TCTTTACCCAAATTTTGTACAAAAG, BP21-R; TTGATCTACAGAGTATGGCAAAAG, BP36-F; GGAAGCTCCTCCAATTAGG, BP36-R; GTAAACCCCTCCTAAACACG, BP38-F; GTGAAAAAGGATTAGAGAAGGG, BP38-R; TTAGTGTGGCACATTTGCC, BP45-F; ATGTTGAATAGATGGATGGATGG, BP45-R; GGAACCGCCTGTCCTTC, BP50-F; AATGGATAGTATCTGTGAATTTTC, BP50-R; AGCTACACAACTTTTTGTATTG.

### EVs labelling & confocal microscopy imaging

HeLa cells were seeded onto glass coverslips pre-coated with poly-D-Lysine for an overnight. Isolated EVs were incubated with 1mM BODIPYTM 493/503 (Final concentration 10 µM; Thermofisher, Cat# D3922) for 20 min at 37°C. Excess dye was removed by centrifuging at 17,000xg for 1h at 4°C and resuspending in PBS. These fluorescently labelled EVs were added to the recipient cells (100 μg/mL) and incubated at 37°C overnight. Then, media was removed, and cells were washed once with PBS. Cells were fixed with 4% paraformaldehyde for 10 minutes and then washed with PBS (3×5min). Cells were permeabilised with Triton-X 100 and washed again. Then, ActinRed™ 555 fluorescent stain (Thermo Fisher Scientific, Catalog# R37112) was added for 30 minutes to label the actin cytoskeleton. Cells were washed with PBS and coverslips were mounted with DAPI mounting media and stored at 4°C until analysis. Confocal Microscope (Leica Stellaris 8) was used to capture images using 63× oil lens from each visual field using the Leica Application Suite Advanced Fluorescence (LAS AF) software (Leica microsystems, Australia) and image analysis was performed using Qupath at default settings (actinRed (red), Bodipy (green), and DAPI (blue)).

### Transmission Electron Microscopy (TEM) of Extracellular Vesicles

For negative staining, carbon-coated 300-mesh copper grids were first glow-discharged in air using a PELCO easiGlow system (20 mA, 30 s, negative polarity) to make the carbon surface hydrophilic. The grids were then placed, carbon-coated side down, onto 5 µL droplets of EV suspension and left for 2 min to allow the vesicles to attach to the grid surface. Excess liquid was carefully removed using Whatman Grade 1 filter paper.

The grids were then washed by placing them onto two separate 5 µL droplets of Milli-Q water, with excess liquid gently removed after each wash. For negative staining, the grids were incubated on a 4 µL droplet of 2% (w/v) aqueous uranyl acetate for 1 min. Excess stain was removed by gentle blotting, and the grids were left to air-dry in the dark before imaging by transmission electron microscopy.

### Oxford Nanopore Sequencing

DNA sample QC was done using Qubit and Genomic DNA screen tape on Tape station to check the quality.

Ligation sequencing (SQK-LSK114) library prep kit was used for Library preparation. 12µL of final library (~230ng) was loaded on PromethION R10 flowcell used for sequencing and ran at high accuracy base calling for 72 hours.

### Predicting ecDNA genomic maps

CoRAL^20^ (version 1.0.r1) was used to reconstruct ecDNA architectures using the long-read whole-genome sequencing data. The samples were sequenced using the Oxford Nanopore (ONT) ligation sequencing kit V14 (SQK-LSK114). The sequencing data was basecalled with high accuracy basecalling (model v4.3.0, 400bps) using Dorado (version 7.4.12). The basecalled data was then aligned to the human reference genome (hg38) based on the wf-alignment workflow (https://github.com/epi2me-labs/wf-alignment). We utilised CNVkit (version 0.9.12) (Talevich et al., 2016) to compute the genome-wide copy number (CN) profile, which was used by CoRAL to identify and filter CN gain regions where amplifications existed. CoRAL was then applied to perform a breakpoint graph construction and cycle decomposition on the CN gain regions to identify candidate ecDNA structures. The outputs of CoRAL were visualised in Circos-style images with CycleViz (version 0.2.0) (https://github.com/AmpliconSuite/CycleViz)^20,21^.

## DATA AVAILABILITY

ONT sequencing data are publicly available in the SRA (BioProject PRJNA1334441).

## ACKNOWLEDGEMENTS

This research was funded by Hearts and Minds Foundation & TDM Growth Partners. Funding to the laboratory of JB is provided by an Investigator Award from the National Health and Medical Research Foundation of Australia (NHMRC APP2026574).

## CONTRIBUTIONS

E.M.P. performed the experiments, contributed to the study design, data collection and interpretation, and manuscript preparation. B.D. performed EV isolation using free-flow electrophoresis (FFE) and characterised the isolated EVs using ZetaView analysis and dot blotting. V.V.P. conducted the bioinformatics analyses. B.Z. assisted with tissue culture experiments, while A.E. contributed to the preparation of experimental materials and stock solutions. J.B. provided access to the FFE technology and equipment and contributed valuable technical expertise and support for the EV separation experiments. O.V. supervised the project. All authors have read and agreed to the published version of the manuscript.

